# Size compensation in Drosophila after generalised cell death

**DOI:** 10.1101/2022.11.04.515194

**Authors:** Noelia Pinal, Natalia Azpiazu, Ginés Morata

## Abstract

Regeneration is a response mechanism to restore tissues that have been damaged or lost by mechanical or physiological insults. We are studying in the wing imaginal disc of *Drosophila* the regenerative response to a high dose of Ionizing Radiation (IR), which we estimate kills at least 35% of the cells. After such treatment irradiated discs are able to restore normal size and shape, indicating there is a mechanism to replace the lost cells. We have tested the role of the Jun N-terminal Kinase (JNK), Janus Kinase (JAK/STAT) and Wingless (Wg) pathways, which have been shown to promote cell proliferation in regenerating tissues. We find that there is size compensation after IR in the absence of function of these pathways, strongly suggesting that they are not necessary for the compensation. We also find that the proliferation rate is not increased after IR. We argue that after generalized death caused by IR there is not a specific mechanism to promote cell proliferation. The irradiated discs suffer a developmental delay and then resume growth at normal rate until they reach the final stereotyped size. The delay appears to be associated with a developmental reversion, as irradiated discs undergo rejuvenation towards an earlier developmental stage. The response to generalized damage is fundamentally different from that to localized damage, which requires activity of JNK and Wg.

## Introduction

Animal tissues react to developmental insults (amputation, injury, irradiation) regenerating the damaged or missing parts through a process that includes compensatory proliferation and reprogramming of the resident cells that survive the damage.

We are studying the mechanisms of regeneration using the *Drosophila* imaginal discs as a model system. These are larval structures that contain the precursor cells of the adult cuticle; they are named after the adult part they differentiate, e.g. eye-antennal disc, wing disc, leg disc and so on. The imaginal discs are classical objects of research in developmental biology; their growth parameters are well known as well as the different genetic factors and signalling molecules involved in their development ^1^. In the case of the wing disc, it begins growth at the interphase between first and second larval instar with about 50 cells to a total of 31000 cells at the end of the larval period^2^. On average the progeny of each initial cell performs 9-10 divisions. It is of interest that the cell division rate is not uniform during the proliferation period; cells proliferate rapidly during the early stages, about 5-6h per cycle, but the rate decreases later in development to about 30h per cycle at late third instar^2^. The disc stops growth at the end of the larval period, at the time pupation starts.

The wing disc has strong regenerative potential; wing disc fragments have been shown to regenerate an entire disc after transplantation in adult hosts^3,4^. More recent experiments^5–7^ also indicate strong *in vivo* regeneration. The common approach in those experiments is to target a particular region of the disc for ablation and then to study how the reminder of the disc reacts and reconstitutes the damaged part. The ablation is achieved by using the Gal4/UAS system to force in the target region expression of pro-apoptotic genes like *hid, eiger* or *reaper*, what generates high apoptosis levels and subsequent cell death. This method allows a precise definition of the target region to ablate. These experiments have identified several factors, like the Jun N-Terminal Kinase (JNK), Wg and Myc pathways, which are implicated in the process^5–8^.

The activity of the JNK pathway appears to be critical for the process. It exerts a dual role: 1-it becomes activated in cells suffering damage, where it triggers apoptosis and subsequent cell death, and 2- JNK-expressing cells secrete mitogenic signals that stimulate the proliferation of neighbour cells – a paracrine function^9^. We have shown that these two functions operate in regenerating wing discs: in an experiment in which wing cells are killed, the moribund JNK-expressing cells release signals that induce additional proliferation of neighbour notum cells, which generate a notum duplicate. If JNK signalling is prevented the duplicated notum does not appear^8^.

Unlike the experiments above in which the damage is restricted to specific disc regions, treatments like ionising irradiation (IR) cause pepper-salt generalised damage, killing a fraction of cells thorough the entire territory of the disc. Published reports indicate that for a dose of 4000R the wing disc can compensate for the loss of 50% or more of the cell population and subsequently attains normal final size^10^. This damage is expected to trigger a generalised response, presumably from cells in the vicinity of the dead ones.

The mechanism(s) by which population size is reconstituted after generalised cell death are unclear. The mode of function of JNK provides an attractive model: as described above, it is activated after IR in cells fated to die but also elicits the secretion of mitogenic signals, which are taken by healthy neighbours; additional divisions performed by the latter could compensate for the cell loss. Some years ago, we showed that two downstream JNK signals, Dpp and Wg, are not involved^11^ in compensatory proliferation, but there is evidence that JNK can activate the JAK/STAT pathway^12,13^ known to induce cell proliferation in imaginal discs (La Fortezza et al., 2016)^14^ and there is also the possibility of other downstream signals.

Moreover, it has been proposed^15^ that in the wing disc much of the size compensation after IR derives from cells of the proximal region of the disc (the hinge), which are refractory to radiation-induced apoptosis: cells from the hinge can migrate to the central region to replace the lost ones. The insensitivity of those cells to apoptosis requires activity of the JAK/STAT and Wg pathways, which implies that both pathways are necessary for the compensation. This hypothesis is supported by lineage data showing that clones from the hinge can extend to the pouch^15^, although it does not explain size compensation in other parts of the disc like the notum.

There are also some experiments suggesting a role of the *p53* gene in the process^16^. These authors found that after IR larvae suffer a significant developmental delay mediated by *p53*, which presumably allows additional time for recovery. However, irradiated *p53* mutant discs reach at the end of the larval period the same stereotyped size as the wildtype, indicating that the mechanism of size compensation does not require *p53* activity.

Thus, there are three pathways that could be involved in the process of size compensation by inducing additional proliferation of cells surviving the IR: JNK, JAK/STAT and Wg. Here we analyse the role of these pathways in irradiated wing discs that lose at least 35% of their cells. We describe their response to IR and also test whether their activity is necessary for compensating for the cell loss. Our principal finding is that irradiated wing discs achieve normal size in the absence of function of JNK, JAK/STAT or Wg. We propose that size compensation after generalised cell death does not require a specific upsurge of compensatory proliferation but that the disc responds to the loss of cells by rejuvenating and prolonging the proliferative period to allow for additional cell divisions until it reaches the final stereotyped size. These results indicate that the response of the wing disc after generalised damage is fundamentally different from that after localised damage

## Results

### Apoptosis and recovery after irradiation in the wing disc

The ability of imaginal discs to recover after IR has been known for a long time. It is illustrated in Figure 1A,B: larvae administered a heavy dose (3000R) of X-rays at second or early third larval period are able to generate mature (prepupal) wing discs indistinguishable from those of non-irradiated larvae. The fact that discs from irradiated and non-irradiated larvae are indistinguishable indicates that the recovery has occurred during the larval period.

**Figure 1.**
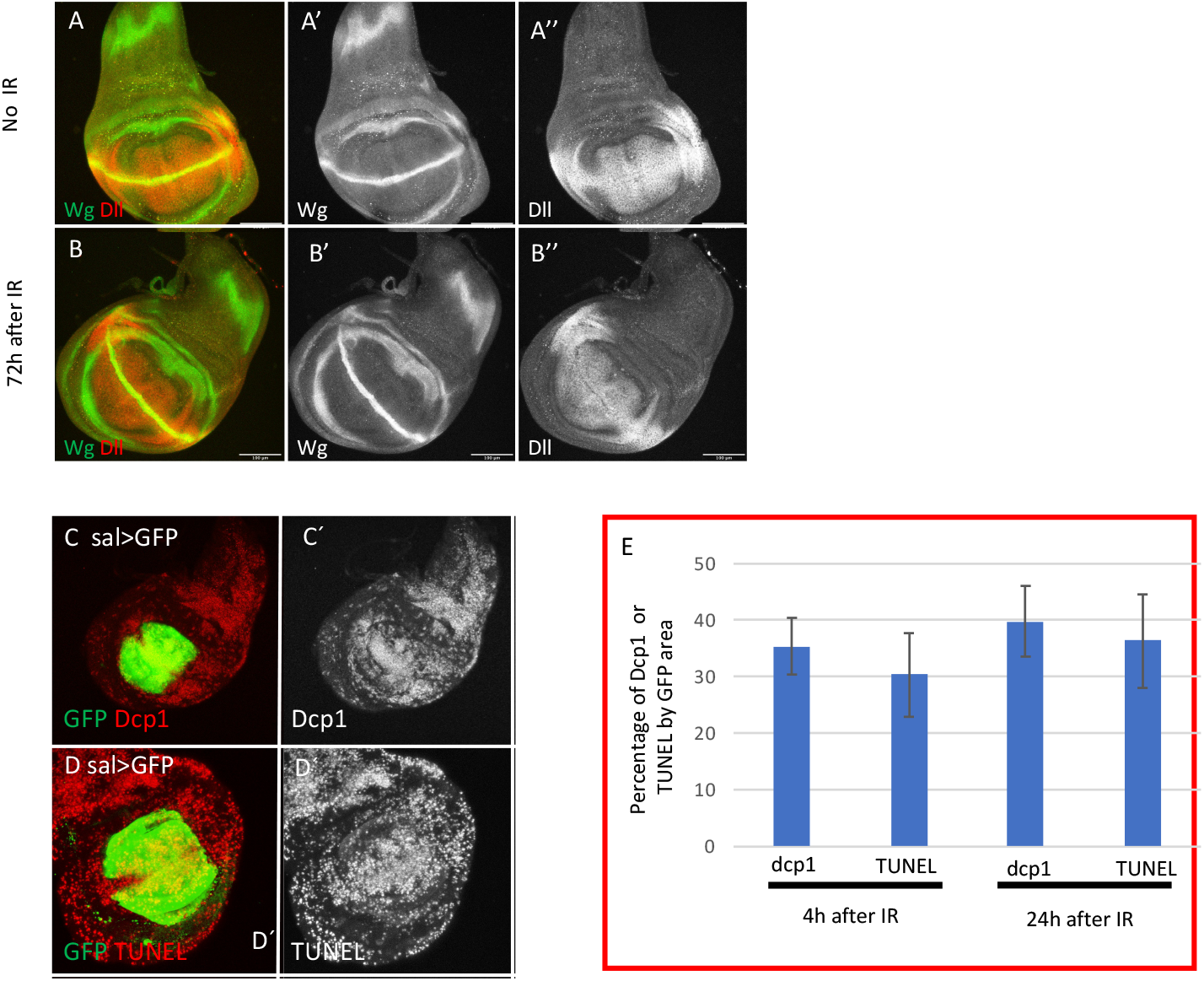
Size restoration and apoptosis after Ionizing irradiation (IR). (A-B) Confocal images of wild type wing imaginal discs of L3 wandering larvae showing Wg (green) and Dll (red) patterns in non-irradiated larvae as control (A-A’’), and in larvae 72h after 3000R (B-B’’). The final size and Wg and Dll expression are fully restored in the irradiated discs. (C-E) Quantification of apoptosis in *sal>GFP* wing discs after irradiation. (C and D) Examples of confocal images of wing discs 24h after irradiation used for the quantification of apoptosis. GFP (green) labels all the cells of the Sal domain, whereas Dcp1 (C-C’) or TUNEL (D-D’) are labelled in red/white. (E) Graph showing the percentage of Dcp1 or TUNEL staining within the GFP area 4h or 24h after irradiation. This value reflects the percentage of cells that are killed by the treatment

To measure the cell lethality caused by IR in our experiments, we irradiated larvae of *sal>GFP* genotype and fixed wing discs 4 and 24h after the treatment. All the cells of the Sal domain (which occupies the central region of the disc, Fig. 1C,D) are labelled with GFP. We then estimated the proportion cells of the domain that contain Caspase activity or TUNEL staining, as indicators of cell death, with respect to those with GFP label. The ratio of cells stained with Dcp1 or TUNEL with respect to those with GFP would be an indication of the amount of the cell death caused by the irradiation. The results are shown in Figure 1E. 4h after the irradiation Dcp1/GFP and TUNEL/GFP ratios are 0.35 and 0.30 respectively, which are increased to 0.39 and 0.36 after 24 h. It is of interest that there are not significant differences in the Dcp1 and TUNEL staining, and since the DNA fragmentation revealed by TUNEL is taken as definitive mark of cell death, it suggests that Dcp1 activity also indicates irreversible apoptosis. We estimate that 3000R cause an overall cell death of about 35%, although this is very likely an underestimate because there is still cell death even 48h after the irradiation^17^.

### Expression of JNK, JAK/STAT and Wg after irradiation

The idea that JNK, JAK/STAT and Wg might be implicated in compensatory proliferation is supported by their response to IR. Up regulation of the JNK and Wg after irradiation have already described in several reports^11,13^: both pathways become de-repressed in regions where they are not normally active. In addition, we find that the expression of JAK/STAT is also modified by IR. It is normally restricted to the hinge region of the disc^18^, but after IR there is also expression in the wing pouch area, clearly visible after 24h (Figure 2). This up regulation of JAK/STAT is not surprising since we have shown that after irradiation it functions downstream of JNK^13^, although it has been reported^19^ that it can function independently of JNK.

**Figure 2.**
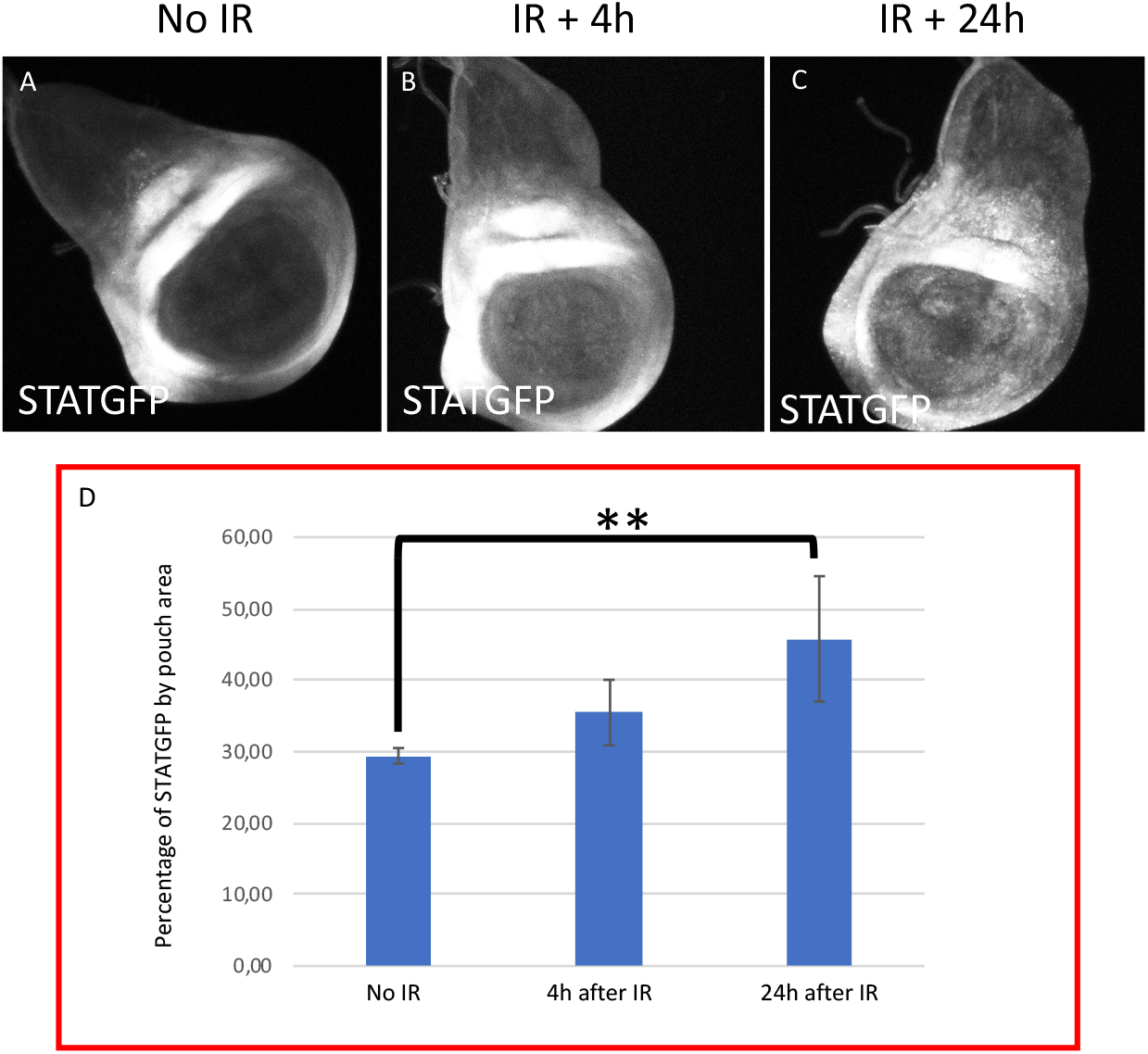
X-radiation induces ectopic JAK/STAT activity. (A-C) Confocal images of wildtype wing discs showing expression of the *STAT-10xGFP* reporter as indicator of JAK/STAT activity. (A) non-irradiated, (B) 4h after irradiation, (C) 24h after irradiation. (D) Graph showing the quantification of STATGFP expression in the pouch region in the control and 4h and 24h after IR. There is significant up regulation of JAK/STAT activity in the wing pouch after IR, which is more pronounced after 24h.

### Rationale of the experiments

Our main goal is to study the response of the wing disc to generalized cell death caused by IR and to identify factors implicated in size compensation. The JNK, JAK/STAT and Wg pathways become ectopically activated after IR (Figure 2) and ^11,13,17^, which suggest that they may play a role in the process.

To assay the implication of the three pathways in size compensation we have made use of the Gal4/UAS Gal80^TS^ system (Fig. 3A). It permits discriminating between the Anterior (A) and Posterior (P) compartments; using the driver line hh-Gal4 we can compromise the activity of the various pathways in the P compartment of irradiated discs, while the A compartment serves as control. If the function of JNK, JAK/STAT or Wg is necessary to compensate for the loss of cells, it would be expected that removing the corresponding function would cause a size reduction in the P compartment of at least 35%. The ratio of the size of the P compartment with respect to the total (P/T), which in normal discs is about 0.4, should be altered significantly.

**Figure 3.**
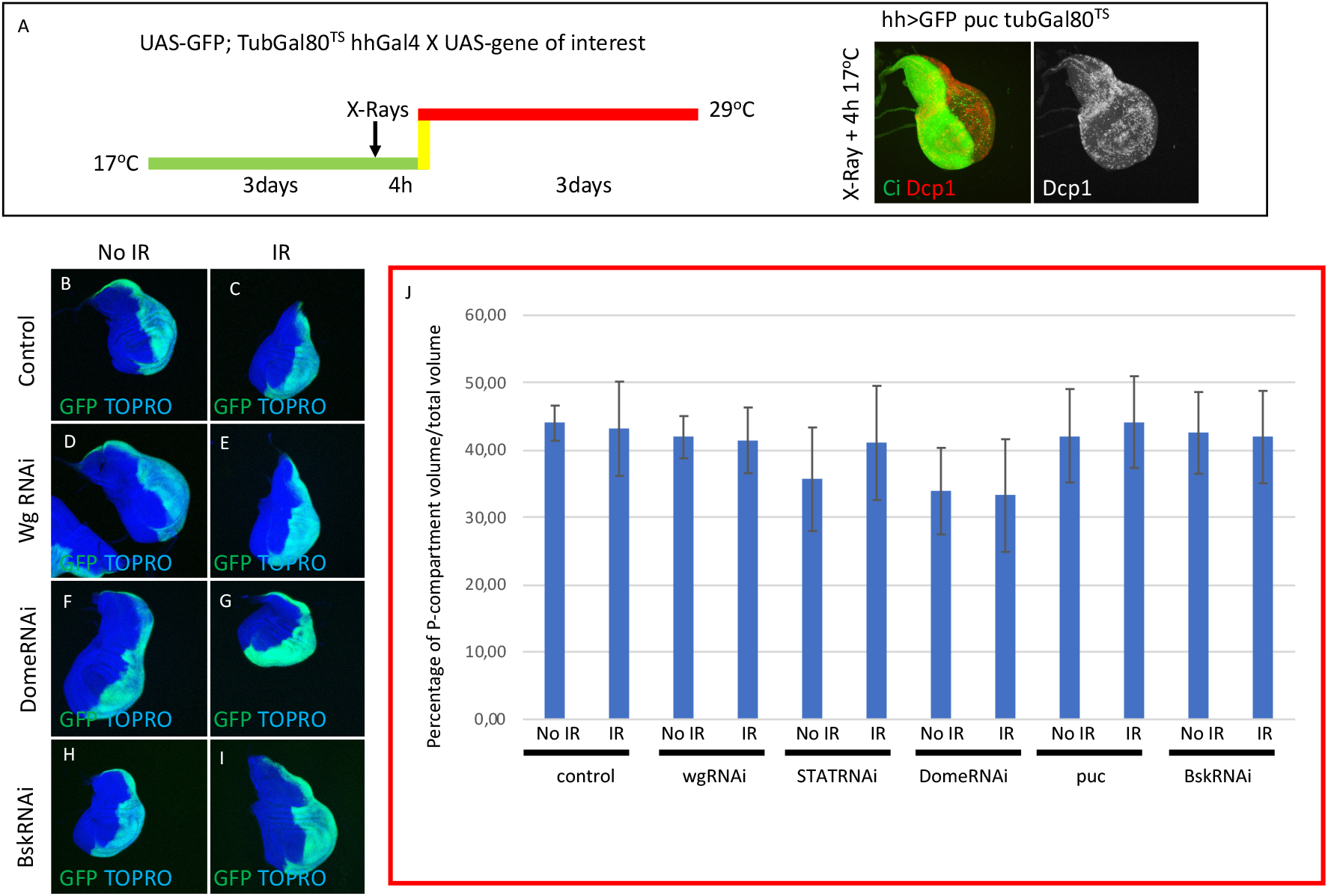
Size compensation in the absence of Wg, JAK/STAT or JNK activity. (A) Schematic representation of the experiment used to assay the role of Wg, JAK/STAT and JNK pathways in size compensation. Larvae of the genotype indicated in the figure were kept at 17°C for 3 days after egg laying before IR. After irradiation they were kept for 4h at 17°C to allow apoptosis and then switched to 29°C to induce the expression of the UAS transgenes that suppress the function of JNK, JAK/STAT or Wg (see Material and Methods for details). The larvae were maintained at 29°C for the rest of development until the wandering stage and then dissected and wing discs discs fixed. The image to the right of A, shows a third instar disc of the genotype indicated stained for apoptosis (Dcp1) 4 h after IR. The anterior compartment is labelled green with the Ci antibody. Note that the amount of apoptosis monitored by Dcp1 (red/white) is the same in the anterior and posterior compartment. (B-I) Examples of confocal images of wing discs used for the quantification, GFP (green) labels the posterior compartment, and TOPRO-3 (blue) the entire disc. Non-irradiated discs on the left column, irradiated ones on the right. (B,C) control discs. (D,E) discs in which Wg pathway is supressed in the posterior compartment by the *UAS-wgRNAi*, (F,G) suppression by the *UAS-DomeRNAi* and (H,I) suppression by *UAS-bskRNAi*. (J) Graph showing the volume of the posterior compartment with respect to the total volume of the disc in the different genotypes studied. There are not significant differences in posterior compartment volume between irradiated and no-irradiated larvae for all the cases. The discs in which the JAK/STAT pathway is suppressed the posterior compartments are smaller, but in both irradiated and no-irradiated discs.

### Suppression of JNK, JAK/STAT and Wg in irradiated P compartments

To suppress the activities of JNK, JAK/STAT and Wg we have used specific RNAi constructs that interfere with the function of the pathways. Suppression of JNK activity can be achieved by RNA interference of the Basket kinase^20^ or overexpressing *puckered*, which functions as a negative regulator of JNK^21,22^. The JAK/STAT pathway can be inactivated by suppressing the function of the transcription factor STAT^15^ or the receptor Domeless^23^, and the Wg pathway by interfering with the ligand itself (see Material and Methods). Those UAS transgenes have been shown to be effective suppressors of the corresponding pathways^8,15^.

Since normal function of JNK is necessary for induction of apoptosis^22^, the analysis of its possible contribution to size compensation requires that JNK be suppressed only *after* cell death has occurred. To address this issue, we have used the thermosensitive dominant Gal4 suppressor Gal80^TS^ to manipulate the time of activity of JNK. Larvae of genotype *UAS-GFP; TubGal80*^*TS*^ *hhGal4; UAS-bskRNAi* (or *UAS-puc*) were irradiated at the second/early third period (3 days at 17°C after egg laying) and shifted to 29°C 4h after IR, thus providing normal JNK function for those 4 h plus the time needed to inactivate the Gal80 suppressor, estimated to be about 6h^24^. We certified that there is apoptosis in those experimental conditions by measuring the levels of Dcp1 4h after IR of third instar discs of genotype *hh>GFP UAS-puc TubGal80*^*TS*^ grown at 17°C and before the shift to 29°C. As illustrated in Figure 3A, 4h after IR at 17°C we find similar levels of apoptosis in the A and P compartments.

To assay the requirements for JAK/STAT or Wg we used a similar protocol (see Material and Methods). The results concerning the requirements of the three pathways for size compensation are shown in Figure 3B-I, J. In comparison with controls (Fig. 3B,C) the size of the P compartment is not reduced after compromising the Wg (Fig. 3D,E), JAK/STAT (Fig. 3F,G) or JNK (Fig. 3 H, I) pathways. The quantified results shown in Fig. 3J indicate that the ratio of P compartment with respect to the total volume of the disc (P/T) is in all cases like in the control non-irradiated discs. In the case of the suppression of JAK/STAT we find that the non-irradiated STAT RNAi or DomeRNAi P compartments are smaller than the normal ones (Figure 3J), suggesting it is required for normal growth of the compartment. However, the relative size of the compartment is not altered by IR. The conclusion is that in the absence of JNK, JAK/STAT or Wg, irradiated discs compensate for the loss of 35% of their cells and attain normal size and shape. These results strongly suggest that these pathways do not play a significant role in size compensation after generalised cell death.

A caveat of the previous interpretation is the possibility of an immediate and strong proliferative response, dependent on JNK or on the two other pathways, within the few hours between the time of irradiation and the suppression of the pathway by Gal80. If that were the case, we would expect a significant augment of cell division markers like EdU in the affected region just after IR. We have tested this hypothesis in an experiment in which we have measured cell division rate by EdU incorporation 5h after IR of discs of genotype *hh>GFP UAS-RHGmiRNA*. The *UAS-RHGmiRNA* construct suppresses the function of *reaper, hid* and *grim*, the three principal pro-apoptotic genes and is a very effective suppressor of apoptosis^13^. As shown in Figure 4A,A’, IR does not cause cell death in the P compartment and hence there is no need for compensatory proliferation, which would be needed in the A compartment. In the hypothesis that there is an upsurge of cell division just after IR we would expect a significant difference of EdU incorporation in the A and P compartments. As illustrated in Figure 4D there is no increase of cell proliferation, monitored by EdU incorporation, neither at 5h nor at 24h after IR (figure 4). These results rule out the possibility of an immediate proliferation response to IR.

**Figure 4.**
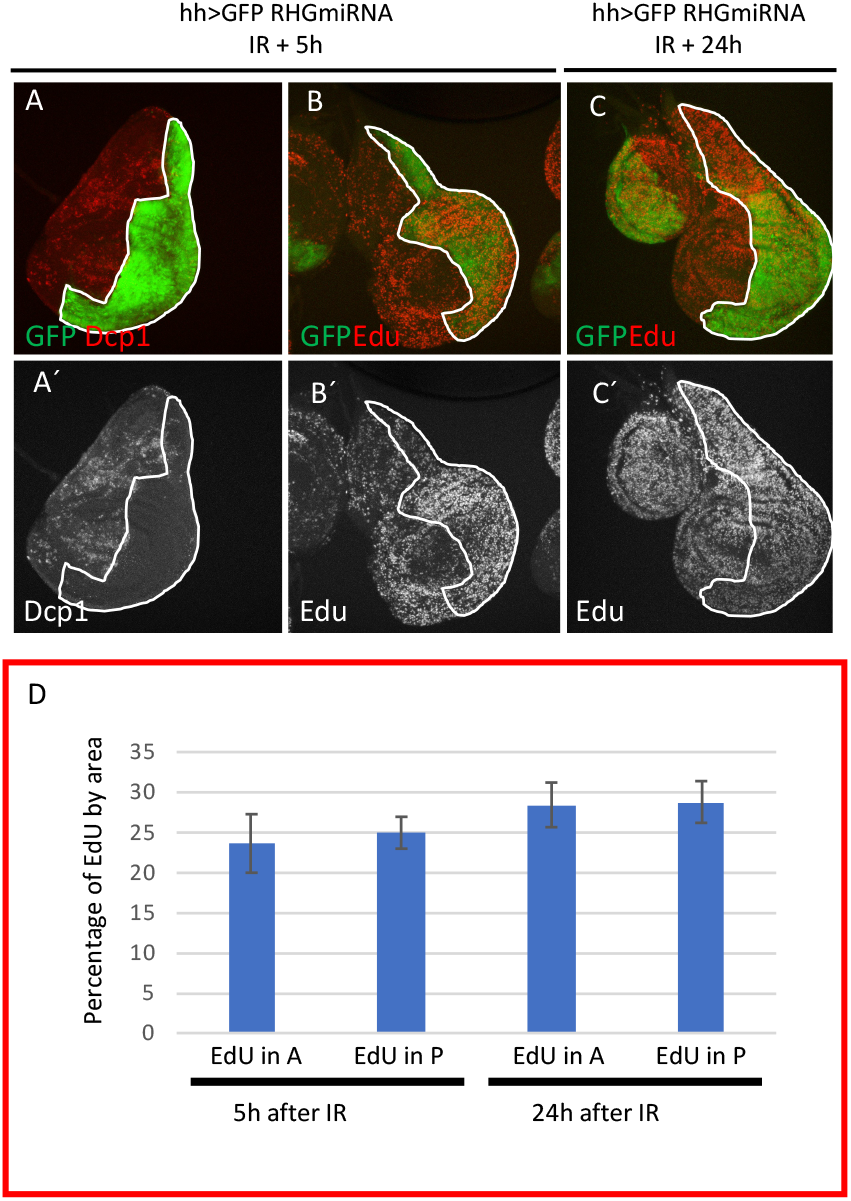
Massive cell loss does not trigger immediate upsurge of cell division after IR. (A-C) Confocal images of wing discs of genotype *hh>GFP RHGmiRNAi*, in which the apoptosis is suppressed in the posterior compartment by the activity of the *UAS-RHGmiRNAi* construct that abolishes the function of the *reaper, hid* and *grim* proapoptotic genes, as illustrated in (A-A’). 4h after IR there are high levels of apoptosis (Dcp1, red) in the anterior compartment but virtually none in the posterior compartment indicating that there is no cell loss. (B-B’) irradiated disc showing equal levels of EdU incorporation (red/white) in the anterior and posterior (labelled with GFP) compartments 4h after the treatment. (C-C’) disc of the same genotype fixed 24 h after the irradiation, also showing similar levels of EdU incorporation. (D) Graph showing quantification of EdU incorporation in the anterior or posterior compartments 5 and 24h after irradiation in discs of the genotype above. It indicates that the loss of cells in the anterior compartment does not induce a upsurge of cell proliferation shortly after irradiation

### Proliferative response after IR

None of the pathways, e.g. JNK, JAK/STAT or Wg, known to be involved in regeneration and growth control appear to be necessary for size compensation after IR. Previous work ^11^ has shown that Dpp is also not involved. It is possible that there may be other factors implicated in generating the additional cell proliferation necessary for the compensation, though it appears unlikely because the former are the major factors involved in the growth, regeneration and patterning of the wing disc.

An alternative is that generalised cell death does not trigger a specific proliferative response. The process for an irradiated disc would be as follows: 1) IR kills a large fraction of the cell population, 2) the affected disc becomes “smaller” than non-irradiated controls and may revert to an earlier developmental stage, 3) the smaller disc resumes growth until it reaches the final stereotype size. This process prolongs the proliferation period of the disc. In essence this model proposes that the affected disc would grow at normal rate, but for a longer time; a temporal regulation.

We have tested this hypothesis by carefully measuring the cell proliferation rates during development of anterior and posterior compartments in discs of genotype *hh>miRHG bsk*^*DN*^. In these discs the cells in the posterior compartment cannot enter apoptosis as they are deficient in the activity of the pro-apoptotic genes. They are also deficient in JNK activity, what prevents the overgrowths caused by the ectopic activation of JNK in apoptosis-deficient cells ^13^. Under these circumstances IR will kill approximately 35% of the cells of the A compartment, whereas there will be no cell death in the P compartment. Using PH3 as marker of cells in mitosis we have measured the mitotic index (MI, number of PH3 dots per surface units in µm^2^) of the Anterior and Posterior compartments of discs 24, 48, 72h after IR.

In 24h discs the MI the A compartment/P compartment ratio is 1.09 (n=16), in 48h discs it is 0.9 (n= 14) and in 72h discs is it 0.6 (n=22). These results indicate that the A and P compartments grow at the same rate for most of the development. The 0.6 ratio of the 72h discs suggests that the P compartment grows faster at the end of development, but since in our experiments this is the compartment in which there is no cell death the proliferation increase cannot be due to size compensation.

### Irradiation reverts the discs to an earlier developmental stage

Previous work^16,25^ has shown that IR causes a developmental delay. As it extends the length of the proliferation period, it may allow for the additional cell divisions necessary to reach the final stereotyped disc size. The slow-down of development in the irradiated discs should be reflected in the expression patterns of genes which evolve during the third larval period, like *wg* ^26^ and *Dll* (describe in this article). We have compared their expression patterns in wing discs from larvae irradiated at 96h AEL and fixed at 120h, with non-irradiated control sibs also fixed at 120h AEL. The results are illustrated in Figure 5; while the control discs show the characteristic patterns of *wg* and *Dll* at mature third instar (Figure 5B-B’’), discs from irradiated larvae are smaller and their *wg* and *Dll* expression patterns correspond to earlier stages (Figure 5 C-C’’). The expression patterns of 96h AEL discs are shown for comparison (Figure A-A’’). In conclusion, irradiated discs remain in a developmental stage earlier than controls. They finally reach mature size (Figure 1B) but it takes additional time, which allow for further cell divisions.

**Figure 5.**
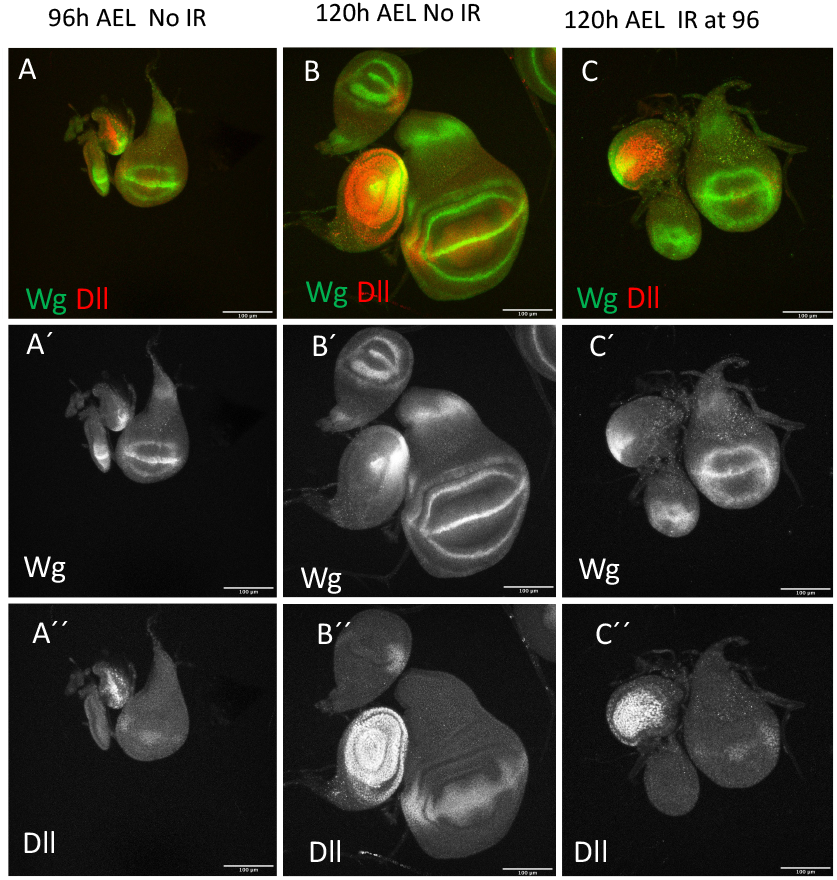
Pause in development after irradiation. (A-C) Confocal images of wing discs of wildtype larvae stained for Wg (green/ white) and Dll (red/white) at different time points of development. (A-A’’) discs of nonirradiated larvae fixed 96h after egg laying (AEL), (B-B’’) discs of non-irradiated larvae fixed 120h AEL. Note that the expression patterns of wg and of Dll are less defined at 96 than at 120 AEL. (C-C’’) discs irradiated at 96h and fixed at 120h AEL. Note that they are smaller than the 120h controls (B-B’’). Also the expression patterns of *wg* and *Dll* are less developed and resemble those of 96h AEL (A-A’’). All scale bars are 100*μ*m.

In an additional experiment, we tested the possibility that as IR makes the discs “smaller”, it could also result in reversion of the irradiated larvae to an earlier developmental stage. After a very short egg laying period of 2h, (to ensure little dispersion in the developmental stage) wing discs from non-irradiated larvae of 96h after egg laying (AEL) were fixed and stained for *wg* and *Dll* expression. In our culture conditions this time correspond to middle third larval period. 96h AEL larvae of the same population as the controls were irradiated at 96h and allowed to develop for further 8 or 24h after IR before wing discs were extracted and fixed.

The expressions of *wg* and *Dll* of controls discs were compared with those of irradiated discs 8h or 24h after IR. The results are illustrated in Figure 6. 8 hours after IR the expression of *wg* is diffuse and not well defined (Figure 6B’), especially along the D/V border. It resembles that described for discs younger that 96h (Martin and Morata, 2006)^26^. Also, the area of *Dll* expression is smaller than that corresponding to 96h discs (Figure 6B’’) and resembles that reported for earlier discs. The discs fixed 24h after IR show (Figure 6C-C’) *wg* and *Dll* patterns similar to those of 96h AEL non-irradiated discs (Figure 6A-A’’). These results strongly suggest that after IR the wing disc reverts to an earlier developmental stage

**Figure 6.**
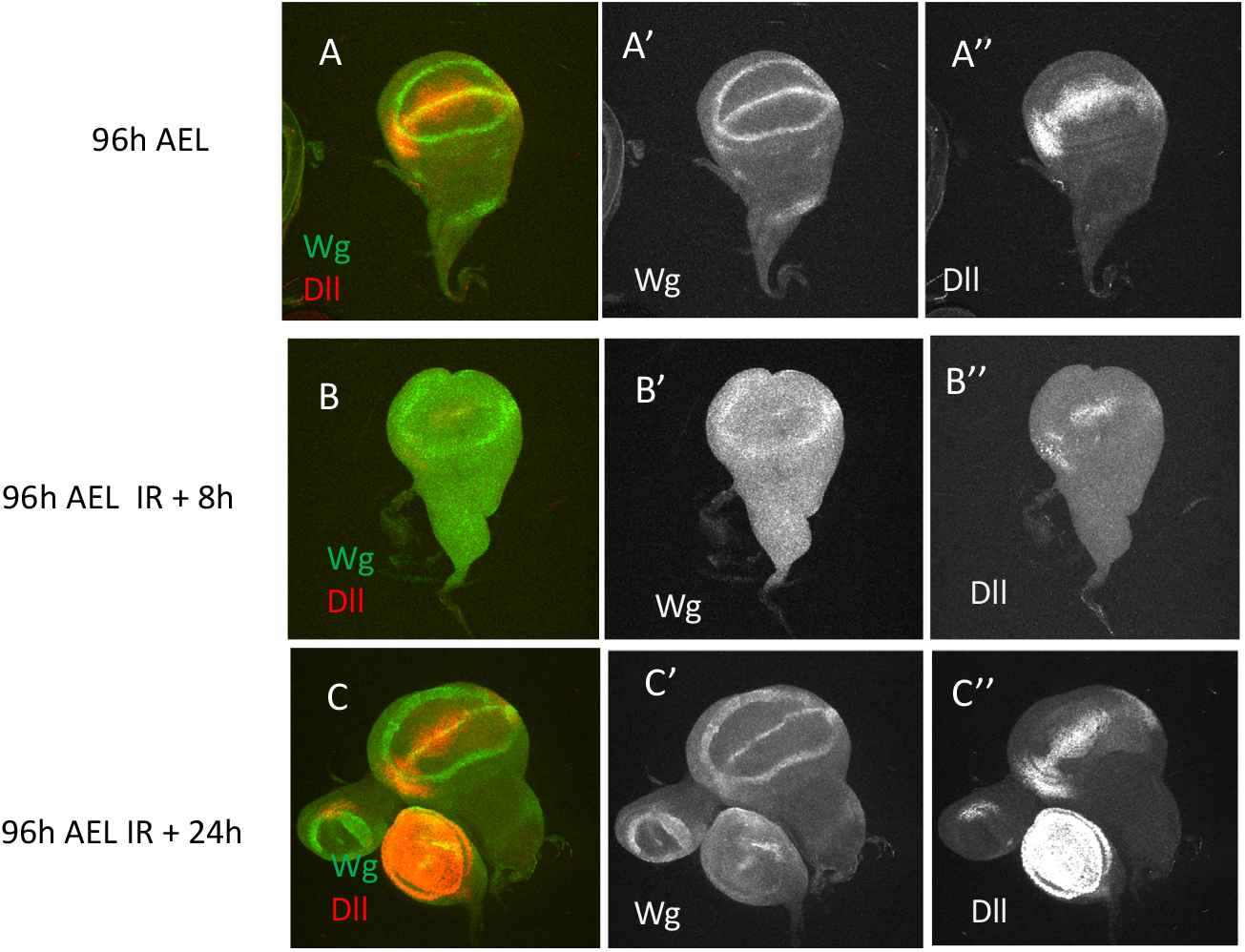
The wing disc becomes rejuvenated after IR. (A-A’’) Canton S wing disc fixed 96h after egg laying (AEL), which corresponds to middle third instar. It has been stained for wg (green or white A’) and Dll (red or white A’’) expression. (B-B’’) Disc of the same genotype irradiated at 96h (AEL) and stained for wg (B’) and Dll expression (B’’) 8 hours after IR. (C-C’’) Disc of the same genotype irradiated at 96h (AEL) and stained for wg and Dll expression 24 hours after IR.

## Discussion

When analysing regeneration process, we can distinguish between the response to localised damage, e.g. amputation or localised injury, with that observed after generalized damage caused by heavy irradiation. The response to localised damage has been analysed in a number of publications^5–8,27^. Key elements in the response are the JNK and Wg pathways, necessary for the additional proliferation required and possibly for the change of identity of the cells involved in the regeneration. It has been shown that during the regeneration process proliferative signalling emanating from dying JNK-expressing cells stimulates the proliferation of neighbour cells^8^.

In the experiments reported here we have analysed the response of the wing disc to IR-induced generalised cell death, which eliminates about 35% of the population. Survivor cells compensate for the lost ones and eventually develop a disc of normal size and shape. Because of its mode of function, the JNK pathway would be expected to be involved in the compensation process; its activation by IR causes cell death by apoptosis, but dying JNK-expressing cells secrete signals that stimulate proliferation of neighbour cells ^9,17,28,11^. This paracrine activity would be responsible for the compensation. Previous work from our laboratory ^11^ showed that there is size compensatory in the absence of the JNK downstream signals Wg and Dpp, but there are other growth factors, directly or indirectly dependant of JNK that could be responsible for the compensation. The experiments reported here rule out a role of JNK in the process as there is size compensation in the absence of JNK function (Figure 3).

Nonetheless, there might be other factors involved in the process. Verghese and Su, 2016 described a proximal region in the wing disc (that they refer to as the “frown”) that contains high expression of wingless and JAK/STAT ^18,26^ and that is refractory to IR-induced apoptosis. The insensibility to apoptosis of the frown cells requires the function of the JAK/STAT and Wg pathways for their absence the amount of apoptosis after IR is like in the rest of the disc. Based on these observations, Verghese and Su, 2016 proposed that the cells from the frown are responsible for repopulating the disc after IR, and also provided some supporting evidence based in cell lineage. The implication of their model is that the property of the frown cells to repopulate the rest of the disc after IR is dependent on JAK/STAT and Wg. However, our experiments demonstrate that there is size compensation in the absence of either Wg or JAK/STAT activity, arguing against the involvement of the cells of the frown.

The *p53* gene has been postulated to play a role in size compensation after IR^16^. IR-treated larvae suffer a slow-down in development, which requires *p53* function. It has been suggested^16^ that *p53* is necessary for the recovery of IR-treated larvae because in *p53* mutant larvae the delay does not occur and the adults that emerge have small wings and multiple morphological abnormalities. However, irradiated *p53* mutant discs attain at the end of larval development the same size as normal discs^16^, indicating that *p53* is not required for size compensation. In *p53* mutants the temporal suppression of apoptosis may allow aneuploidy and other damaged cells to survive, which may be responsible for the abnormalities observed in IR-treated *p53* mutant flies.

The general conclusion from all the above is that none of the factors examined, JNK, JAK/STAT, Wg, p53, is responsible for the size compensation after IR. This is in striking contrast with the response observed after localised damage.

### There is no compensatory proliferation after generalised damage

Our results indicate that in the wing disc, and by extension in the other discs, the mechanism to restore size and shape after generalised damage is very different from that operating after localised damage or amputation. A localized damage like the ablation of the wing pouch or the entire wing region causes up regulation of the JNK pathway^5–8^, which plays a major role in the process. Its activity is associated with the death of damaged cells, but also causes secretion of proliferative signals necessary for the resident cells to regenerate the missing or damaged tissue ^8^. Thus, there is JNK-dependant compensatory proliferation.

In contrast, after generalised damage like that caused by IR, the activities of pathways associated with regeneration like JNK, Wg or JAK/STAT do not have a role in the compensation process. These pathways are indeed activated after IR ^11,13^ and Figure 2, likely a reflection of their intrinsic response to damage of any kind and probably caused by the production of Reactive Oxygen Species in the damaged tissue^29^. However, their activities are inconsequential regarding size compensation: in their absence the disc restores normal size after the death of at least 35% of the cells.

In the IR-treated *hh>miRHG bsk*^*DN*^ discs we have measured the cell proliferation rate in the anterior compartment, in which there is large amount of cell death, and in the posterior compartment, in which cell death is impeded. We find no difference in the mitotic index of these compartments. Thus there is no need for a specific mechanism to induce additional cell proliferation of resident cells after IR damage. Strictly speaking there is no compensatory proliferation in the sense of a special wave of cell divisions triggered by the damage.

In our view the key factor in the restoration process would be the mechanism that controls overall size, which stops growth once the disc has attained the stereotyped final size. After IR the discs lose a fraction of the cells, which makes them smaller (although the alteration of size is not easy to visualize) and presumably as a consequence the irradiated discs revert to an earlier developmental stage (Figure 6).

We propose that generalised cell death, like that caused by IR, does not trigger a proliferative response; it simply rejuvenates the affected discs, which revert to an earlier developmental stage. It prolongs the length of the proliferation period, what allows the surviving cells to perform additional divisions at normal rate until the disc reaches its final stereotyped size. Size compensation after generalised damage is a temporal mechanism.

## Materials and Methods

### Drosophila strains

The *Drosophila* stocks used in this study were: *hh-Gal4, sal*^*Epv*^*-Gal4* (gift from J. F. de Celis, CBMSO, Madrid, Spain), *tub-Gal80*^*ts*^, *UAS-RHGmiRNA, STAT-10xGFP, UAS-BskRNAi* (5680R1, National Institute of Genetics stock center, Japan), *UAS-WgRNAi, UASBskRNAi, UAS-STATRNAi* (33637), *UAS-*DomeRNA*i* (32860), *UAS-GFP, UAS-puc*^*2A*^ and canton wild type (Bloomington Drosophila Stock Center).

### Tissue staining

Immunostaining was performed as described previously^30^. Images were captured with a Leica (Solms, Germany) DB5500 B confocal microscope. The following primary antibodies were used: rabbit anti-Dcp1 (Cell Signalling, antibody #9578) 1:200; mouse anti-Wingless (DSHB 4D4) 1:50; guinea pig anti-Dll (gift of Dr Carlos Estella^31^); rat anti-Ci antibody (DSHB 2A1) 1:50. Fluorescently labelled secondary antibodies (Molecular Probes Alexa) were used in a 1:200 dilution. TO-PRO3 (Invitrogen) was used in a 1:600 dilution to label nuclei. All the n number stated in the text represent individual larvae of the mentioned genotypes. All experiments are replicated at least three times.

### Dcp1/TUNEL and STATGFP quantifications

To estimate the percentage of cells that undergo apoptosis after irradiation larvae of the genotype *sal*^*Epv*^*-Gal4/UAS-GFP* were irradiated with a 3000R dose 4 or 5 days after egg laying. Irradiation was performed with an X-Ray machine Phillips MG102. Wing imaginal discs from these larvae were stained with Dcp1 antibody or TUNEL 4h or 24h after irradiation. Confocal images were processed with FIJI software to measure the areas covered by GFP, Dcp1 and TUNEL labels. With these data the percentage of Dcp1 or TUNEL areas within the Sal domain was obtained.

A similar procedure was used to quantify the modification of JAK/STAT activity after irradiation, using the *STAT-10xGFP*^15,24^ reporter. Larvae bearing the reporter were irradiated 4 or 5 days after egg laying, and the wing discs fixed 4h or 24h after irradiation. Confocal images from these discs were processed with FIJI software to measure in the pouch region (normally devoid of JAK/STAT activity) the gain of function of JAK/STAT, which is not normally active in that area. The comparison of irradiated discs and non-irradiated controls allows to estimate the gain of JAK/STAT activity after X-rays.

### Pathway inhibition in posterior compartment and quantifications

To block Wg, JAK/STAT or JNK activity in posterior compartments after irradiation, we designed a cross in which males bearing one of the transgenes *UAS-WgRNAi, UAS-DomeRNAi, UAS-STATRNAi, UAS-BskRNAi* or *UAS-puc* were crossed to *UAS-GFP; tub-Gal80*^*ts*^, *hh-Gal4* females. Larvae originating from these crosses were raised at 17°C and irradiated with a 3000R dose 3 days after egg laying. Irradiation was performed with an X-Ray machine Phillips MG102. Afterwards the larvae were kept at 17°C for 4h to allow apoptosis in normal genetic conditions and then shifted to 29°C for 3 days before dissection. Non-irradiated controls were treated with the same procedure except the irradiation. Wing imaginal discs were stained with TO-PRO3. Confocal images were processed with FIJI voxel counter plugin software to measure total disc volume (TO-PRO3 staining) and posterior compartment volume (GFP staining).

### EdU incorporation and quantification

Larvae of the genotype *UAS-RHGmiRNA/UAS-GFP; tub-Gal80*^*ts*^, *hh-Gal4* were raised at 25°C for 3 days, then shifted to 29°C to allow expression *RHGmiRNA*, which impedes apoptosis in the posterior compartment. 24h after the temperature shift larvae were irradiated with a 3000R dose. Wing imaginal discs were dissected 5h and 24h after irradiation for EdU labelling. Wing imaginal discs were cultured in 1 mL of EdU labelling solution for 30 min at room temperature and subsequently fixed in 4% paraformaldehide for 30 min at room temperature. Rabbit anti-GFP (Invitrogen) 1:200 antibody was used overnight at 4°C before EdU detection to protect GFP fluorescence. EdU detection was performed according to the manufacturer instructions (Click-iT EdU Alexa Fluor 555 Imaging Kit, ThermoFisher Scientific). TO-PRO3 was used to label the entire disc area.

To quantify EdU incorporation in the anterior and posterior compartments confocal images were processed with the FIJI software to measure total disc area (TO-PRO3 staining) and posterior compartment area (GFP staining) and EdU staining in each of these areas. Anterior area and EdU staining were obtained by the difference between total and posterior. With these data the percentage of EdU incorporation by area (anterior or posterior compartment) was obtained.

### Expression patterns of Wg and Dll after irradiation

Eggs from a canton wild type stock were collected in agar plates supplemented with sugar, juice and propionic acid, with a 4h interval. Next morning larvae were collected every hour for 7h and place in two/three new fly food tubes. First collection of the day was discarded. Larvae were kept at 25°C for 3 days. At this time point wing imaginal disc were dissected from one of the tubes collected every hour to stain with anti-Wg and anti-Dll antibodies. The larvae from the other tube were irradiated with a 3000R dose. 24h after irradiation wing discs from irradiated and non-irradiated larvae were dissected and stained with anti-Wg and anti-Dll antibodies. Finally, 3 days after irradiation wing imaginal discs from irradiated and non-irradiated larvae were dissected and stained with anti-Wg and anti-Dll antibodies.

## Acknowledgements

We thank the members of our lab for comments and discussions and Angélica Cantarero and Rosa Gonzalez for technical help. We also thank Ernesto Sánchez-Herrero for comments and suggestions. This work has been supported by grants PGC2018-095151-B-100 of the Ministerio de Ciencia, Innovación y Universidades, FEDER funds and the Agencia Estatal de Investigación and CSIC intramural funding 2021EP007.

